# Rehabilitating the benefits of gene tree correction in the presence of incomplete lineage sorting

**DOI:** 10.1101/2025.07.09.663893

**Authors:** Manuel Lafond, Celine Scornavacca

## Abstract

Gene trees play an important role in various areas of phylogenomics. However, their reconstruction often relies on limited-length sequences and may not account for complex evolutionary events, such as gene duplications, losses, or incomplete lineage sorting (ILS), which are not modeled by standard phylogenetic methods. To address these challenges, it is common to first infer gene trees using fast algorithms for conventional models, then refine them through species tree-aware correction methods. Recently, it has been argued that such corrections can lead to overfitting and force gene trees to resemble the species tree, thereby obscuring genuine gene-level variation caused by ILS. In this paper, we challenge and refute this hypothesis, and we demonstrate that, when applied carefully, correction methods can offer significant benefits, even in the presence of ILS.

## 1 Introduction

Gene trees are fundamental components in several downstream phylogenomics analyses. Deciphering the evolutionary histories of gene families provides insights into how genes diversify and adapt, and has applications ranging from the prediction of genes with similar functions, to enhancing our understanding of the processes of gain and loss and of the mechanisms that can lead to the expansion of gene families. Naturally, the accuracy of such biological inferences depends heavily on the accuracy of the gene trees themselves.

This contrasts with the fact that gene trees are difficult to infer with precision. Gene sequences are typically of limited length and thus convey only limited evolutionary signals [26, 23]. Furthermore, genes undergo evolutionary events that are not incorporated into typical phylogenetic reconstruction models. In particular, gene duplication events followed by divergent evolutionary fates can lead to heterogeneous mutation rates across the tree, with gene losses complicating the picture [18, 6, 10].

These challenges can be overcome by first reconstructing gene trees using standard models, then correcting them while accounting for such events. A common strategy is to collapse low-support branches of a gene tree and resolve the resulting polytomies using information from the species tree. For instance, the tool TRACTION finds a refinement of the gene tree with minimum Robinson-Foulds distance to the species tree [2], whereas ecceTERA and profileNJ find the refinement with a minimum duplication and loss cost [8, 16] (see also [29, 30]). TreeFix uses a different approach and explores the space of trees around the input one, to find one with a smaller duplication and loss cost but a comparable likelihood [25]. Such correction methods were shown useful in joint inference of gene trees and species trees [1], detecting whole genome duplications [4], or predicting orthology and paralogy relationships [11], for example.

These correction algorithms only consider duplications, losses, and horizontal gene transfers (HGT, which exchange genes between co-existing species) even though incomplete lineage sorting (ILS) is also known to hinder the reconstruction process. Under ILS, alleles may persist through speciation events, producing gene trees that diverge from the species tree even in the absence of duplications or losses [3, 13, 28]. While there are methods that take ILS into account [12, 17], they are often computationally intensive and impractical at the genome scale, where thousands of gene families are analyzed. For instance, StarBEAST2 [17], MrBayes [20], and BPP [5] are Bayesian approaches that were shown to be very accurate in the presence of ILS, but are quite slow compared to IQ-TREE, a maximum likelihood (ML) phylogenetic inference tool [15]. In [27], it is reported that the latter always takes less than 33 seconds to infer 100 gene trees on their datasets, whereas StarBEAST2 and MrBayes require up to 2 hours and 37.5 hours, respectively. Other statistical approaches such as PHYLDOG [1] can accurately perform joint reconstruction of gene and species trees, but suffer from the same scalability issues. This prevents these accurate tools from scaling up to run on larger datasets and, for that reason, are rarely used in phylogenomics studies where many gene trees are involved. Instead, fast but possibly less accurate methods such as ML phylogenetic inference tools, which do not account either for HGT or ILS, are much more popular in practice. This motivates the need for quick correction procedures that can address gene tree estimation errors of ML tools while retaining efficiency.

However, Yan et al. [27], recently made a case against gene tree correction in the presence of ILS. One of the main arguments against current methods for correction is that ILS results in gene and species tree discordance, and thus that correcting the gene tree without accounting for ILS will make it resemble the species tree and inevitably produce errors. To support this, they perform simulations to compare TreeFix and TRACTION against the uncorrected predictions from IQ-TREE, and their results demonstrate that the two correction approaches often increase gene tree estimation error.

First, we note here that the tool ecceTERA was not evaluated in Yan et al.’s study, despite showing very good performances for non-ILS data sets [9]. Also, TreeFix, unlike several other correction algorithms, had already been shown to over-fit the data [21], so we do not consider it as representative method. Moreover, a single bootstrap support threshold of 75% was evaluated, which is arguably high and can lead to heavy modifications of the gene trees. Thus we decided to re-evaluate the dataset from [27] and demonstrate that gene tree correction, when applied judiciously, does improve accuracy even in the presence of ILS.

## Results

We first summarize the previous work and proceed with the results of our re-analysis.

### Previous simulations, runs and findings

In Yan et al. [27], a species tree *S* with 11 taxa (including one outgroup) was simulated using TreeSim [24]. The branch lengths were rescaled to make the root of the ingroup of height of 2, 5, and 10 in coalescent units, respectively, corresponding to high, medium, and low ILS rates (shorter branches are more subject to ILS). Then, coalescent simulations were performed along *S* to generate datasets composed of 100 gene trees (each with 10 replicates and a single individual per species, with no duplications nor losses). Seq-Gen [19] was used on each gene tree to simulate a multiple sequence alignment. The authors tested two mutation rates *θ* ∈ {0.001, 0.01 }and three possible number of sites in {200, 800, 2000 }. IQ-TREE was used to reconstruct the gene trees with 100 bootstrap replicates. This produced 120 datasets of 100 gene trees, and each gene tree was corrected with TreeFix and TRACTION. For the latter, branches with bootstrap support below 75% were contracted. As a measure of gene tree estimation error, the uncorrected and corrected gene trees were compared to the true simulated tree using the unrooted Robinson-Foulds distance.

To summarize the findings of [27] on these datasets, TreeFix generally improves accuracy on short sequences of length 200, has mixed results on sequences of length 800, and has decreased accuracy on longer sequences of length 2000, especially on higher mutation rates *θ* = 0.01. As for TRACTION, in most cases its accuracy is close but slightly worse than the uncorrected gene trees, with a more pronounced decrease under high levels of ILS. See [27, Figure 2] for detailed results.

### Our runs and findings

For rehabilitating the benefits of gene tree correction, we choose to rerun tests using ecceTERA, a program that implements a fast parsimony reconciliation algorithm accounting for gene duplication, gene loss and HGTs [8]. Here, we reconstruct gene trees using IQ-TREE and use the correction feature of ecceTERA, in which branches with a bootstrap support lower than the given threshold are collapsed by ecceTERA, and rearranged by finding a refinement with a minimum duplication and loss cost, or duplication, loss and transfer cost [9] (we used the ecceTERA default costs for each event).

We corrected gene trees under two different settings: the first considers only duplications and losses as the macro-evolutionary events influencing gene histories, while the second additionally incorporates gene transfer events. All other ecceTERA parameters are set to their default values. For both settings, we run ecceTERA with three different support thresholds: 50, 70 and 90 (in the Appendix, we also include an analysis with thresholds between 10 and 40, but they do not give better accuracy).

We then computed the normalized unrooted Robinson-Foulds distance between the simulated gene trees and the ones reconstructed with IQ-TREE, and between the simulated gene trees and the IQ-TREE trees corrected by ecceTERA. Figure 1 shows the result when correcting using duplications and losses, while Figure 2 also considers HGTs.

**Figure 1.**
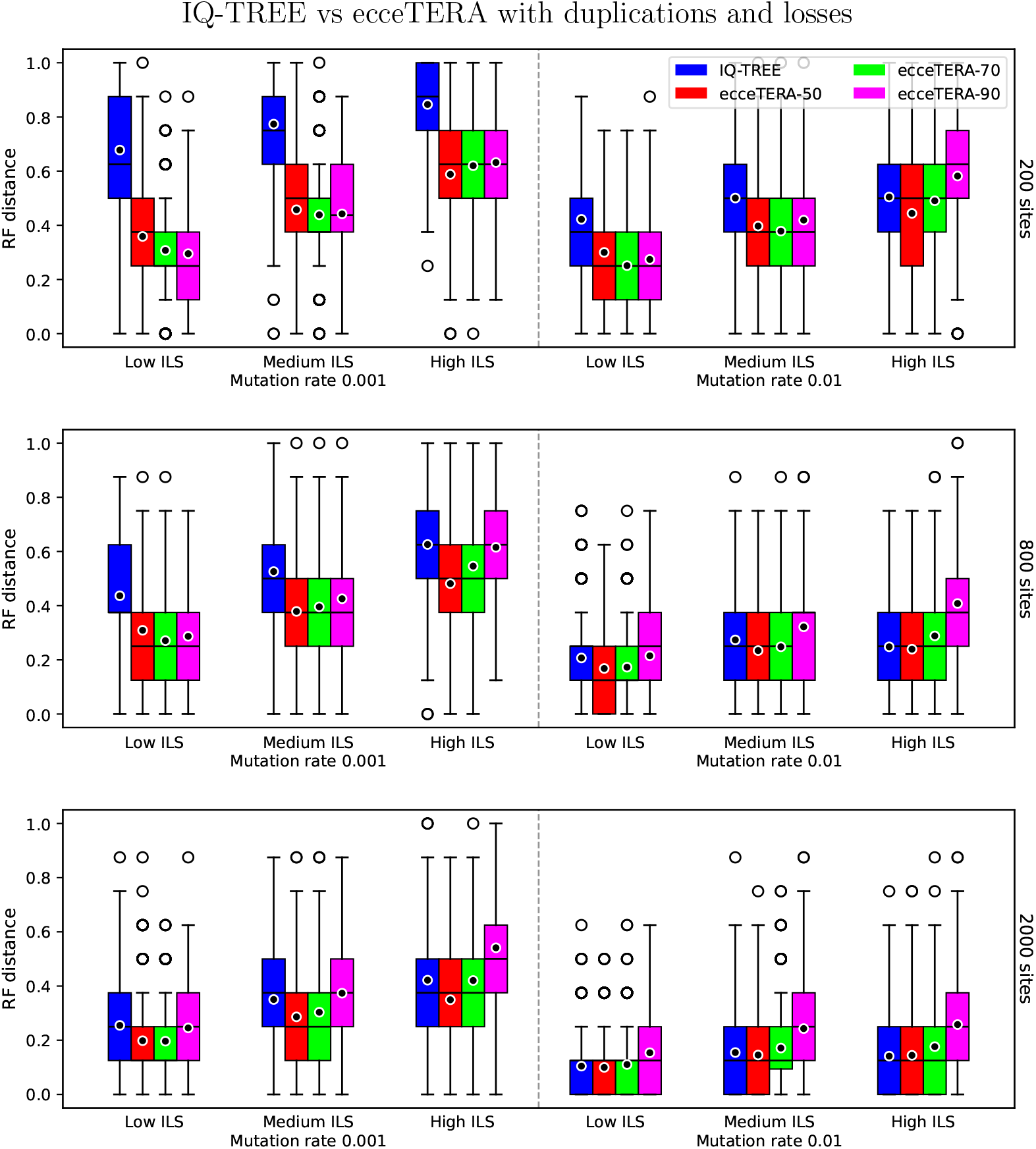
Gene tree error estimation (via the normalized unrooted Robinson-Foulds distance) of IQ-TREE and ecceTERA under thresholds 50, 70, and 90, under the DL model. Black circles represent mean distances.

**Figure 2.**
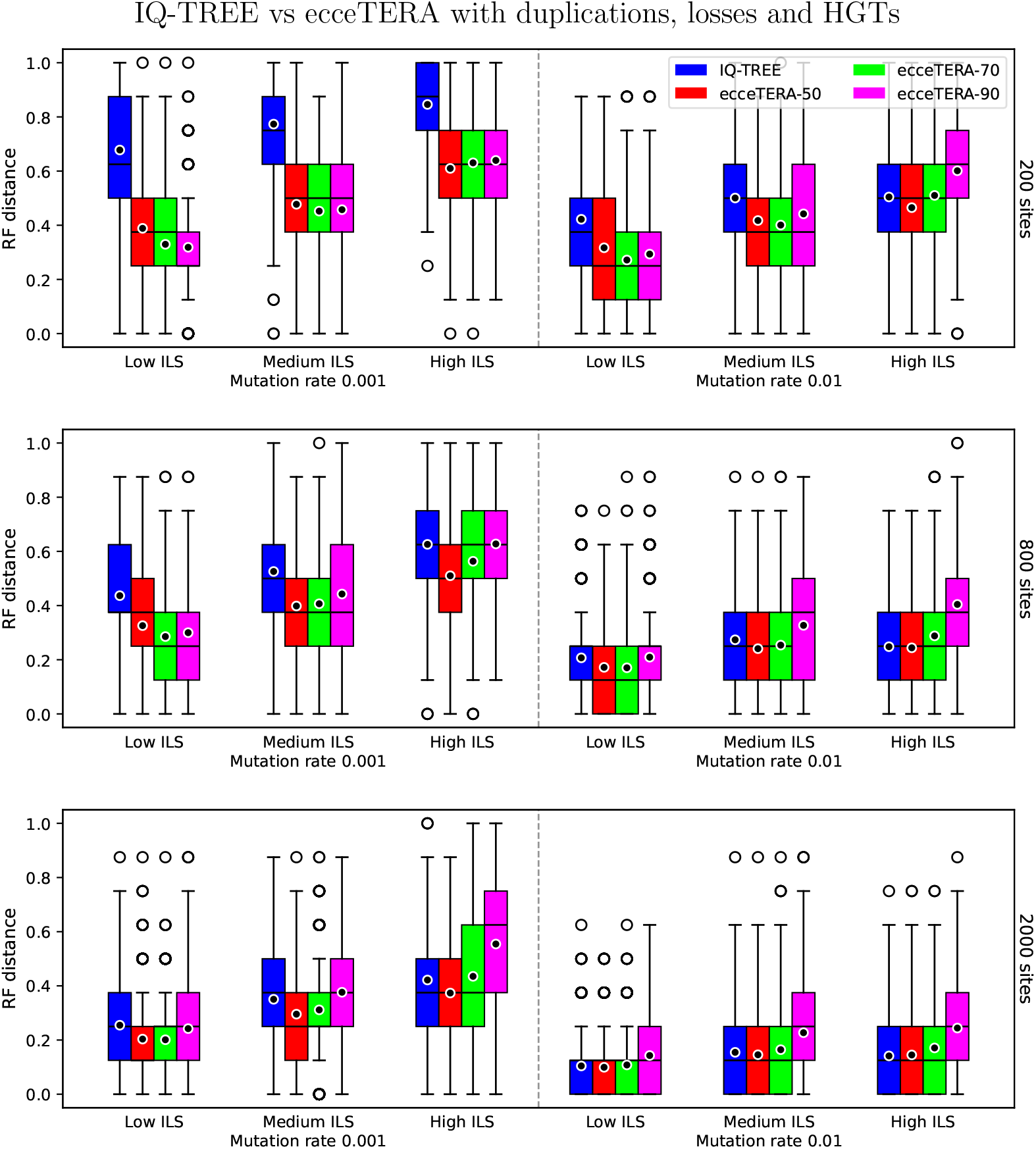
Gene tree error estimation (via the normalized unrooted Robinson-Foulds distance) of IQ-TREE and ecceTERA under thresholds 50, 70, and 90 under the DTL model.

**Figure 3.**
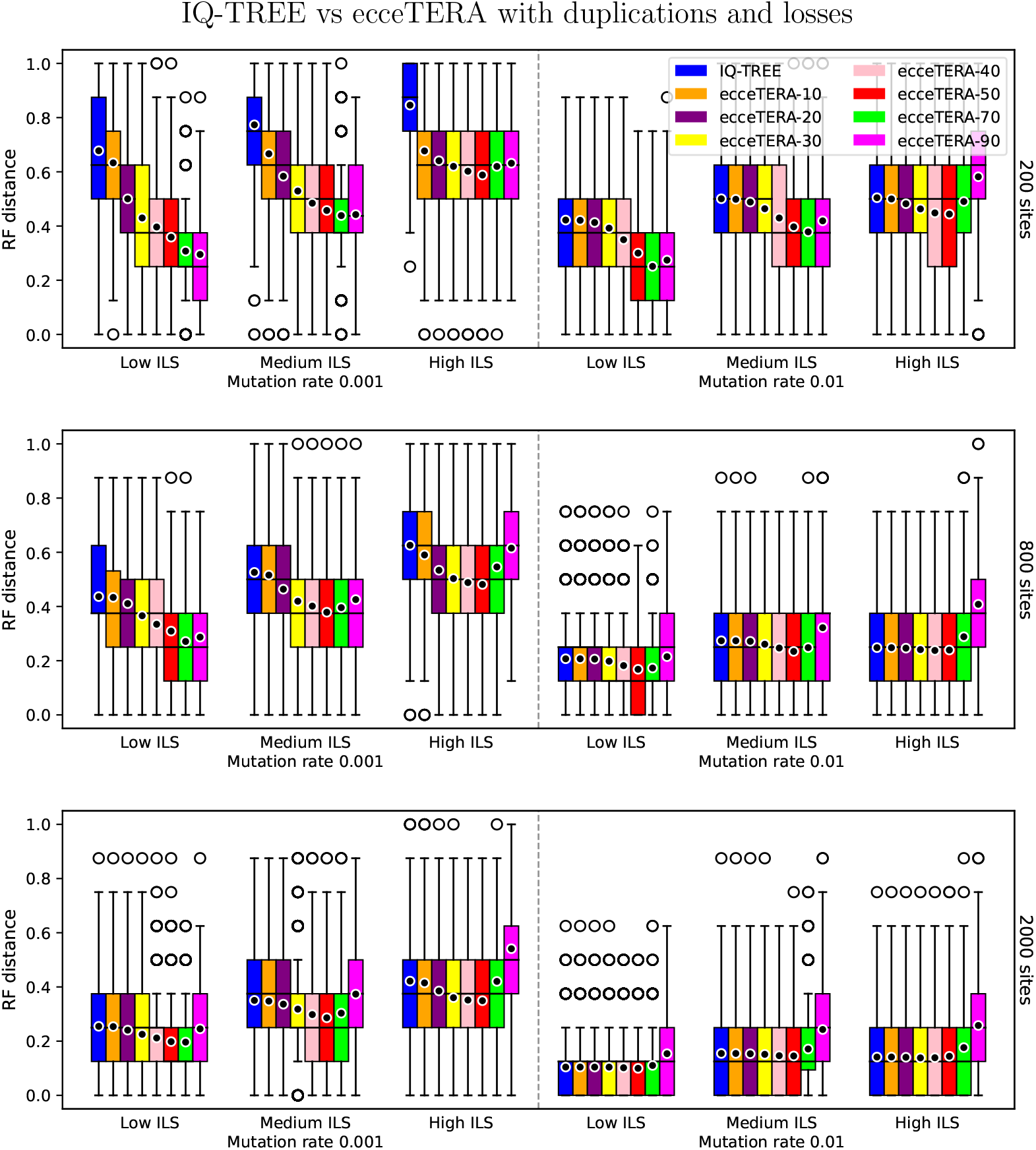
Box-plot of normalized unrooted RF distances for each threshold under the DL model.

**Figure 4.**
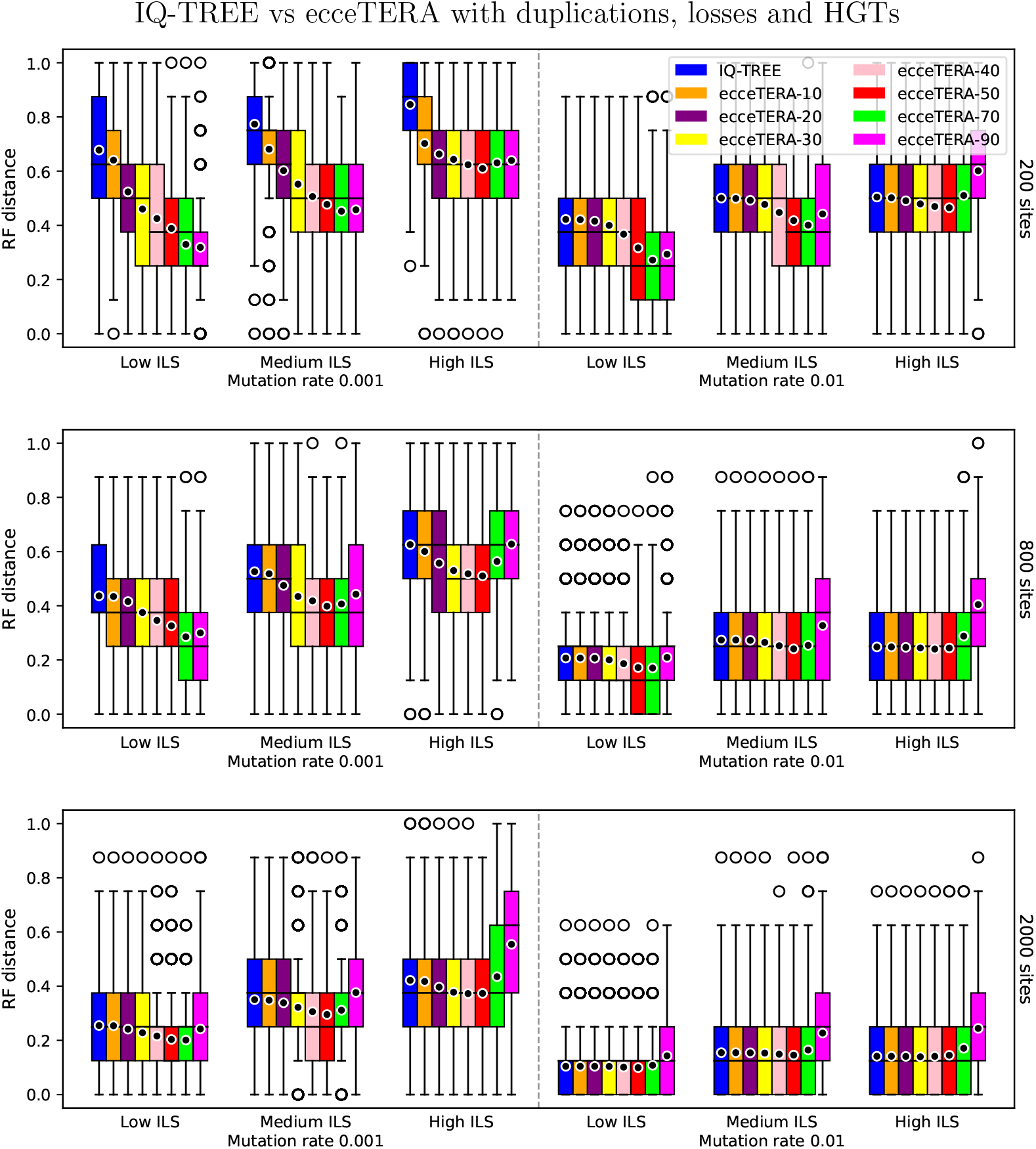
Box-plot of normalized unrooted RF distances for each threshold under the DTL model.

Looking at Figure 1, the initial observation to highlight is that implementing corrections generally proves beneficial across all parameters examined. The box plots using threshold 50 exhibit a lower or equal estimation error box than the uncorrected version, being strictly better in most cases. This becomes particularly noticeable for the combination of a mutation rate of 0.001 and 200 sites, where reconstruction errors tend to occur more frequently and where correction has more impact.

Additionally, using a threshold of 50 generally appears to be more beneficial than the higher thresholds, with exceptions occurring with 200 sites and low/medium ILS, and 800 sites and low ILS. Moreover, while the advantages of correction are more pronounced when ILS is low, the benefits of correction are also noticeable under high ILS, contrasting with the hypothesis proposed in [27] that correction frequently increases errors.

In Figure 2, we can see that correcting taking into account gene transfer is still beneficial compared to the IQ-TREE reconstructions, even if less pronouncedly. In almost all cases, the mean error is slightly higher than in Figure 1 (see for example with 200 sites and medium ILS, the median is higher for threshold 90 on mutation rate 0.001, and on mutation rate 0.01 the box is extended upwards). Nonetheless, the error on threshold 50 corrections is very similar to the duplication and loss corrections, and still generally better than the uncorrected trees. For those cases, additionally considering gene transfer as possible macro-evolutionary events seems to add too much liberty to the correction of gene topologies, leading to overfitting.

## Discussion

The results shown in the previous section contrast with the findings of Yan et al. We believe that this is due to several key factors.

First, while it is true that branches of gene trees discordant with the species tree may result from ILS and should not always be corrected, such regions are often well-supported by bootstrap analyses. A conservative threshold thus preserves these branches. On the other hand, poorly supported branches may reflect reconstruction errors which methods such as ecceTERA are specifically designed to correct. The analysis in [27] was limited to a support threshold of 75%, which probably explains how this was overlooked. Our results, including the data on lower thresholds shown in the Appendix, suggest that a bootstrap of 50% is safe as it does not make the trees worse, while offering good error reduction potential. The threshold echoes earlier studies [7, among others], which advocated using the majority-rule criterion in phylogenetics. Second, the methods chosen by the authors may not be the most appropriate for gene tree correction. TRACTION optimizes the Robinson-Foulds distance, which is not directly related to evolutionary events that affect genes (although it may build accurate species trees from gene trees). TreeFix, while more sophisticated, explores a vast tree space and may overfit gene trees to the species tree, as shown in [21]. In contrast, methods such as ecceTERA address both concerns by explicitly modeling duplications and losses and only modifying poorly supported regions.

Let us also note here that the dataset that we re-analyzed contains gene trees that evolve exclusively under ILS, without duplications, HGTs, or losses. While this is sufficient for our goals here, that is, to rehabilitate correction methods when ILS is present, future experiments should focus on gene trees that also include these events in addition to ILS. In this case, the gene trees are probably much more difficult to reconstruct using sequence-based methods, and we expect correction methods to provide even more meaningful improvements. Future experiments should also question the impact of ILS on gene and species tree co-estimation methods such as AleRax [14].

While we agree that caution is required when correcting gene trees in the presence of ILS, our results show that applying correction methods appropriately has clear benefits. Rather than discarding these approaches, they should be used with care to improve the reliability of phylogenomic inferences, even when ILS is present in the data.

## Acknowledgments

A preprint version of this article has been peer-reviewed and recommended by PCI Evol Biol (https://doi.org/10.24072/pci.evolbiol.100872).

## Data availability

All data and code needed to reproduce the experiments are available at https://zenodo.org/records/17653545 [22].

## Funding

ML acknowledges the Natural Sciences and Engineering Research Council (NSERC, grant RGPIN-2019-05817) for the financial support of this project.

## Conflict of interest disclosure

The authors have no conflict of interest to declare.

## Appendix

Below we include the results for all considered bootstrap thresholds.

## Notes

### Competing Interest Statement

The authors have declared no competing interest.

### Summary of Updates

PCI badge and reference to the fact that the preprint has been peer-reviewed and recommended by PCI Evol Biol

